# A Rapid Bead-Based Assay For Screening Of SARS-CoV-2 Neutralising Antibodies

**DOI:** 10.1101/2021.10.13.464050

**Authors:** Santhik Subhasingh Lupitha, Pramod Darvin, Aneesh Chandrasekharan, Shanakara Narayanan Varadarajan, Soumya Jaya Divakaran, Sreekumar Easwaran, Shijulal Nelson-Sathi, Perunthottathu K Umasankar, Sara Jones, Iype Joseph, M. Radhakrishna Pillai, TR Santhoshkumar

**Author notes:** Equal contribution (Santhik Subhasingh Lupitha, Pramod Darvin, Aneesh Chandrasekharan).

## Abstract

Quantitative determination of neutralizing antibodies against Severe Acute Respiratory Syndrome Corona Virus-2 (SARS-CoV-2) is paramount in immunodiagnostics, vaccine efficacy testing, and immune response profiling among the vaccinated population. Cost-effective, rapid, easy-to-perform assays are essential to support the vaccine development process and immunosurveillance studies. Here, we describe a bead-based screening assay for S1-neutralization using recombinant fluorescent proteins of hACE2 and SARS-CoV2-S1, immobilized on solid beads employing nanobodies /metal-affinity tags. Nanobody-mediated capture of SARS-CoV-2 - Spike (S1) on agarose beads served as the trap for soluble recombinant ACE2-GFPSpark, inhibited by neutralizing antibody. The first approach demonstrates single-color fluorescent imaging of ACE2–GFPspark binding to His-tagged S1-Receptor Binding Domain (RBD-His) immobilized beads. The second approach is dual-color imaging of soluble ACE2-GFPSpark to S1-Orange Fluorescent Protein (S1-OFPSpark) beads. Both methods showed a good correlation with the gold standard pseudovirion assay and can be adapted to any fluorescent platforms for screening. Life-time imaging of the ACE2-GFPSpark confirmed the interaction of ACE2 and S1-OFPSpark on beads. The self-renewable source of secreted recombinant proteins from stable cells and its direct use without necessitating purification renders the platform a cost-effective and rapid one than the popular pseudovirion assay and live virus-based assays. Any laboratory with minimum expertise can rapidly perform this bead assay for neutralizing antibody detection using stable engineered cells.

## Introduction

Coronavirus disease 2019 (COVID-19), caused by the Severe Acute Respiratory Syndrome Corona Virus-2 (SARS-CoV-2), has resulted in extensive morbidity and severe fatality worldwide. Aggressive drug discovery and vaccine development efforts are being pursued by academia and pharmaceutical industries targeting major viral proteins. Large-scale vaccination programs are already ongoing in several countries employing a wide variety of vaccines (Kim, Marks, & Clemens, 2021). Currently, it is not clear how many of them will emerge as successful vaccines in the long run in controlling the pandemic. The well-studied and confirmed entry mechanism of the virus is mediated by the Receptor Binding Domain (RBD) of viral spike S1 protein that interacts with the host cell receptor, Angiotensin-I converting enzyme-2 (ACE2) (Tai et al., 2020). This vital entry pathway remains the golden target for any drug intervention or vaccine development approaches (Huang, Yang, Xu, Xu, & Liu, 2020).

Suitable, easy-to-perform, and cost-effective assays are required to study the interaction of SARS-Cov-2 S1-protein with the ACE2 receptor for drug discovery and to evaluate the neutralization efficacy of antibodies produced in response to either natural infection or a vaccine. Even though S1 protein reactive antibody detection using ELISA, lateral flow devices, and Luminex assays have been developed and available as *in vitro* diagnostic tools and adapted for the quantitative determination of antibody titer (Carter et al., 2020), they do not precisely determine the neutralization activity of antibody in preventing S1 protein-ACE2 interaction. Virus neutralization is validated via classical virus neutralization tests using live virus infection in permissive cells carried out in a BSL3 laboratory or pseudovirion assay in a BSL2 laboratory (Yang et al., 2020). Considering the urgent requirement of potential vaccines and drugs, diverse, innovative models were described using the latter approach. Pseudovirion assays involve transfection of cells with plasmid vectors encoding SARS CoV-2 spike protein and packaging proteins of heterologous viruses. Multiple versions of Vesicular stomatitis virus (VSV) pseudotyped neutralization assays were described (Schmidt et al., 2020). Similarly, Lentiviral pseudotyping using 2^nd^ generation or 3^rd^ generation system is also being described (Dautzenberg, Rabelink, & Hoeben, 2021; Witting, Vallanda, & Gamble, 2013). Recently, M.Eugenia *et al*. described a replication-competent VSV system, rVSV-SARS-CoV-2 S, to identify spike-specific therapeutics and viral entry inhibitors (Dieterle et al., 2020). Although the pseudovirion assay remains the best approach to screen neutralizing antibodies, it requires time-consuming, complex transfections with multiple plasmids to generate pseudovirions and subsequent reinfection in permissive cell models. Also, it is to be noted that most of the S protein pseudotyping performed in other virus models is not as efficient as that of VSVΔG pseudotyping and is labor and time-intensive assays. Better cell-free or cell-based systems that allow rapid and easy testing of large libraries or vaccine candidates are in high demand. A newer ELISA-based platform was described recently to detect anti-SARS-CoV-2 neutralizing antibodies blocking HRP-conjugated RBD protein from binding to the hACE2 protein pre-coated on an ELISA plate (Tan et al., 2020). This assay also requires costly reagents and time to perform the assay. A promising bioluminescence-based biosensor to detect the binding of SARS-CoV-2 viral spike (S) protein to its receptor, angiotensin-converting enzyme 2 (ACE2), and its utility for drug screening and neutralizing antibody testing was also described (Azad et al., 2021).

We describe a new bead-based system to screen ACE2 -S1 protein binding inhibitors using fluorescent-tagged recombinant proteins and nanobodies. Among these, the nanobody-based approach using agarose beads immobilized with S1-OFP and recombinant soluble ACE2-GFP performed well in most image-based detection platforms. Also, we generated cells expressing S1-OFP and ACE2-GFP that served as the self-renewable source of these recombinant proteins that can be directly used without protein purification steps, rendering the assay simple, user-friendly, cost-effective, and rapid. Further, we substantiated the assay’s utility for neutralizing antibody screening using a panel of validated neutralizing antibodies and the result correlated with pseudovirion readout. The assay is easy to adapt to diverse image-based platforms such as automated microscopy, well-plate imaging, and Highthroughput (HTS) systems and is expected to support the performance evaluation of the ongoing global vaccination and studies to understand the neutralizing antibody dynamics.

## Materials and Methods

### Cells and expression vectors

HEK293T cells and SiHa cells were procured from the central cell line repository of RGCB (CCL RGCB). CaCo-2 cells were obtained from ATCC. DLD-1 cells were obtained from Sigma (Merck #90102540). The cells were routinely maintained in DMEM (Thermo Fisher Scientific #61965026) supplemented with 10 % Fetal Bovine Serum (Thermo Fisher Scientific #26140).

Codon optimized SARS-CoV-2 (2019-nCoV) Spike S1 Gene with C terminal OFP Spark fluorescent tag (pCMV3-C-OFPSpark) was obtained from Sino biological (#VG40591-ACR). Full-length DNA clone of Human angiotensin I converting enzyme (peptidyl-dipeptidase A) 2 with C terminal GFPSpark tag (pCMV3-hACE2-GFPSpark) was obtained from Sino biological (#HG10108-ACG). The mammalian expression plasmid for RBD of SARS-CoV-2 protein S (spike) with 8xHis tag was obtained from Addgene (#145145). The mammalian expression vector for soluble ACE2 pcDNA3-sACE2 (WT)-sfGFP (#145171) was obtained from Addgene. The expression vector anti-RBD Sybody (Addgene #153522) was a kind gift from Prof. Markus A. Seeger. GFP nanobody and mCherry nanobody prokaryotic expression vectors were procured from Addgene (#49172, #70696). The transfection grade plasmids were prepared using Qiagen Midi Kit (#12143). The RFP Trap beads were procured from Chromotek (#rta10). The expression vector used for the pseudovirion SARS CoV2-19 was from Addgene (pCDNA 3.1_Spike_Del19 #155297). The packaging vectors and Lenti ZsGreen were from Addgene (pHIV-Zsgreen #18121, psPAX2 #12260).

### Generation of high expressing ACE2-GFP and S1-OFP stable cells

HEK293T cells were transfected with ACE2-GFPSpark (Sino Biologicals) using FuGENE HD transfection reagent (Promega #E2311) and maintained in Hygromycin B (Sigma Aldrich #H3274) for two weeks. Cells were sorted based on green fluorescence intensity and grew in 6 well plates in low density. Multiple clones expressing varying levels of ACE2-GFPSpark were expanded and employed for the study. Similarly, HEK293T cells were transfected with S1-OFPSpark, and multiples clones with variable expression levels were expanded. To generate dual stable cells, HEK293T cells expressing moderate level of both ACE2-GFP and S1-OFP were developed employing flow sorting and colony selection. Eukaryotic expression vector RBD-Venus was created by the Vector Builder Inc (Chicago, IL, USA).

### Affinity purification of proteins

Lysates prepared from the transfected cells were processed for purification using respective affinity tags as per standard protocol. We expressed and purified anti-GFP and anti-mCherry nanobodies fused to GST in *E. coli*. Cell lysates containing GFP and OFP fusion proteins were pulled down with GST-tagged anti-GFP and anti-mCherry nanobodies pre-bound to glutathione– Sepharose-4B beads (GE Healthcare #17075601), respectively, as per the standard protocol.

### Western blot

Cells were lysed in lysis buffer containing 1% NP-40 and protease inhibitor cocktail by incubating in ice for 30 min. Proteins were estimated by using Bradford reagent, and 40 μg of proteins were resolved electrophoretically on 8–15% SDS polyacrylamide gels, transferred to nitrocellulose membranes, and detected by chemiluminescence (Amersham, Buckinghamshire, UK # GERPN2209). Antibodies used and their dilutions are given in table 1.

### Immobilization of GFP Nanobody to Agarose beads

The bacterial expression vector for anti-GFP nanobody pGEX-6P-1 was procured from Addgene. *E. coli* cells were transformed with the expression vector followed by induction using 0.1 mM IPTG (Sigma Aldrich #I6758) for 4 h at 30°C. The pelleted cells were lysed and used to purify the recombinant proteins with glutathione–Sepharose 4B beads (Sigma Aldrich #GE17-0756-01). The purified protein or whole cell lysate was incubated with the respective agarose beads at various time points with continuous agitation and washed before imaging. Nonspecific binding was inhibited by preincubating the beads with 3% BSA.

### High-throughput imaging approach

Two different high throughput imaging platforms were employed in the study. The custom-configured microscope includes an automated microscope Nikon Ti equipped with Lumencore light solid-state light engine and Andor EMCCD Camera ixon895. The beads, after proper staining, were seeded onto the 96-well glass-bottom plates imaged using 10x 0.45 NA objective or 20x 0.75 NA objective. The images were acquired and analyzed using NIS-Elements software.

### Mouse immunization and serum preparation

BALB/c mice were immunized with purified RBD-His protein expressed in *E. coli* as per the standard protocol approved by IAEC.

### Life-time imaging of ACE2 GFP – S1-OFP interaction on RFP-Trap beads

Fluorescent life-time imaging experiments were performed on SP8 FALCON (FAst Lifetime CONtrast), an advanced confocal-based microscope for Time-Correlated Single Photon Counting (TCSPC) from Leica Microsystems, Germany. This equipment is configured with a white light pulsed laser with a spectral range of 470 to 670 nm and an Acousto-Optical Beam Splitter (AOBS) for filterless emission tuning and detection using sensitive GaAsP detectors. The beads were placed on an 8-well glass-bottom chamber (Nunc) and imaged using a dedicated 100x oil immersion objective with a numerical aperture of 1.4. Life-time data analysis was done using built-in algorithms and represented as a histogram. For confocal imaging, ACE2 GFPSpark and Spike OFPSpark were excited at 488 nm, and 540nm laser lines and emission of 490 -530 and 545 to 570 nm were collected in sequential mode. For life-time measurements, only donor was excited, and life-time was recorded with the same excitation-emission band.

### Confocal Imaging

The cells or beads were placed on a 96-well imaging plate and imaged under 20x 0.75 NA objective using Nikon Confocal Imager A1R equipped with all-solid-state lasers from Agilent. For real-time imaging of S1-OFP binding, cells were maintained in an on-stage incubation chamber and were excited with a 562 nm laser, and red emission was collected at an interval of 5 minutes up to 1 hour.

### SARS-CoV-2 Pseudovirion Assay

HEK293T cells were stably expressed with humanACE2 tagged with HA following transfection and selection. A lentiviral-based pseudotyping was employed using lenti-ZsGreen. HEK293T cells were transfected with lenti-ZsGreen (Addgene) and 10 µg of psPAX2 (Addgene) and SARS-CoV-2-S variant (pCDNA 3.1_Spike_Del19, Addgene) at a ratio of 1:2:1 using the transfection reagent (Lipofectamine 3000, Thermo Fisher Scientific #L3000001). Six hours after the transfection, the medium was replaced with fresh medium and allowed to grow for 72 hours. The supernatant containing the pseudovirion particles was collected, centrifuged, filtered, and concentrated using PEG 8000 and used immediately at appropriate MOI or stored at −80°C. For the neutralization assay, HEK293T cells stably expressing HA-ACE2 were seeded on 96 well optical bottom plates and grown up to 70% confluence. The antibody at indicated dilutions in DMEM was pre-incubated with an equal volume of SARS-CoV-2 pseudovirions for 2 hours for inhibition. The mixture was added to the HEK293T-HA-ACE2 cells for 4 hours, followed by fresh medium, and wells with IgG served as control. The wells were imaged under a fluorescent microscope for GFP fluorescence, and the percentage of positive cells was calculated and compared with IgG and SARS CoV2 pseudovirion only well.

## RESULTS AND DISCUSSION

### Bead-based Single color fluorescence imaging of ACE2 binding to SARS-CoV-2-S1 protein

The entry of SARS-CoV-2 to the host cell is mediated by viral S1 protein binding to the receptor hACE2. Most of the host ACE2 positive cell types are targeted by the virus, and their functional impairment accelerates the disease progression in a subset of patients with underlying morbidities (Wu, Deng, Li, & Yang, 2021). The RBD domain of the S1 protein that mediates the viral entry is highly antigenic and elicits a good immune response (Tai et al., 2020). Remarkable seroconversion also has been demonstrated against Spike protein in COVID 19 patients that are in line with the decreased overall mortality rate among affected people and provides hopes for the disease management with the ongoing vaccination efforts (Wolff et al., 2020).

The popular ELISA-based antibody detection approach cannot distinguish the neutralizing activity of an antibody response. The SARS-CoV-2 virus neutralization test and pseudovirion assays used for neutralizing antibody testing are time-consuming and labor-intensive assays. Better rapid assay platforms are necessary for large-scale screening of neutralizing antibodies. To develop a rapid assay for testing the antibody’s neutralizing activity to block the binding of S protein to ACE2, we have utilized cost-effective agarose affinity beads as the binding platform. SARS-CoV-2-RBD-His expressed either in *E. coli* or HEK293T stable cells served as an RBD-His source, which binds to ACE2. The recombinant RBD domain of S1 protein with His-tag was purified using an affinity column (Figure 1 a). The purified protein was immobilized on agarose beads (Ni-NTA) using Histidine tag as shown in the schematic diagram to be used as a trap for soluble ACE2 (Figure 1 b). Similarly, we have expressed the human ACE2 as a green fluorescent fusion protein in HEK293T cells, and whole-cell extract of the stable cells served as the source for ACE2-GFPSpark (referred to as ACE2-GFP from here on) (Figure 1 c & d). The RBD-His agarose beads showed a time-dependent increase in binding of ACE2-GFP when used at a concentration of 100 μg/ml diluted whole cell lysate (Figure 1 e) with an average bead density of 100 beads per ml.

**Figure 1.**
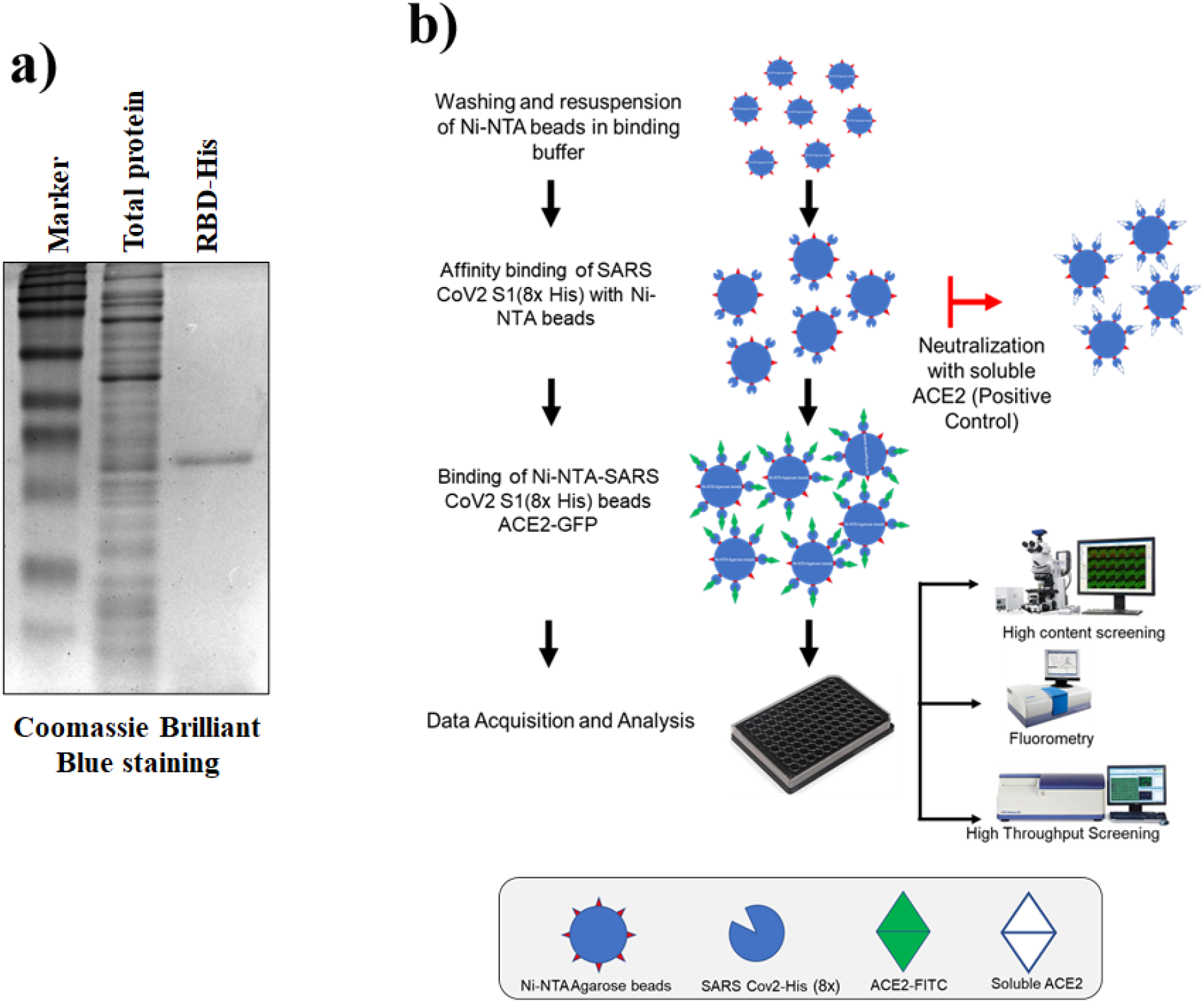
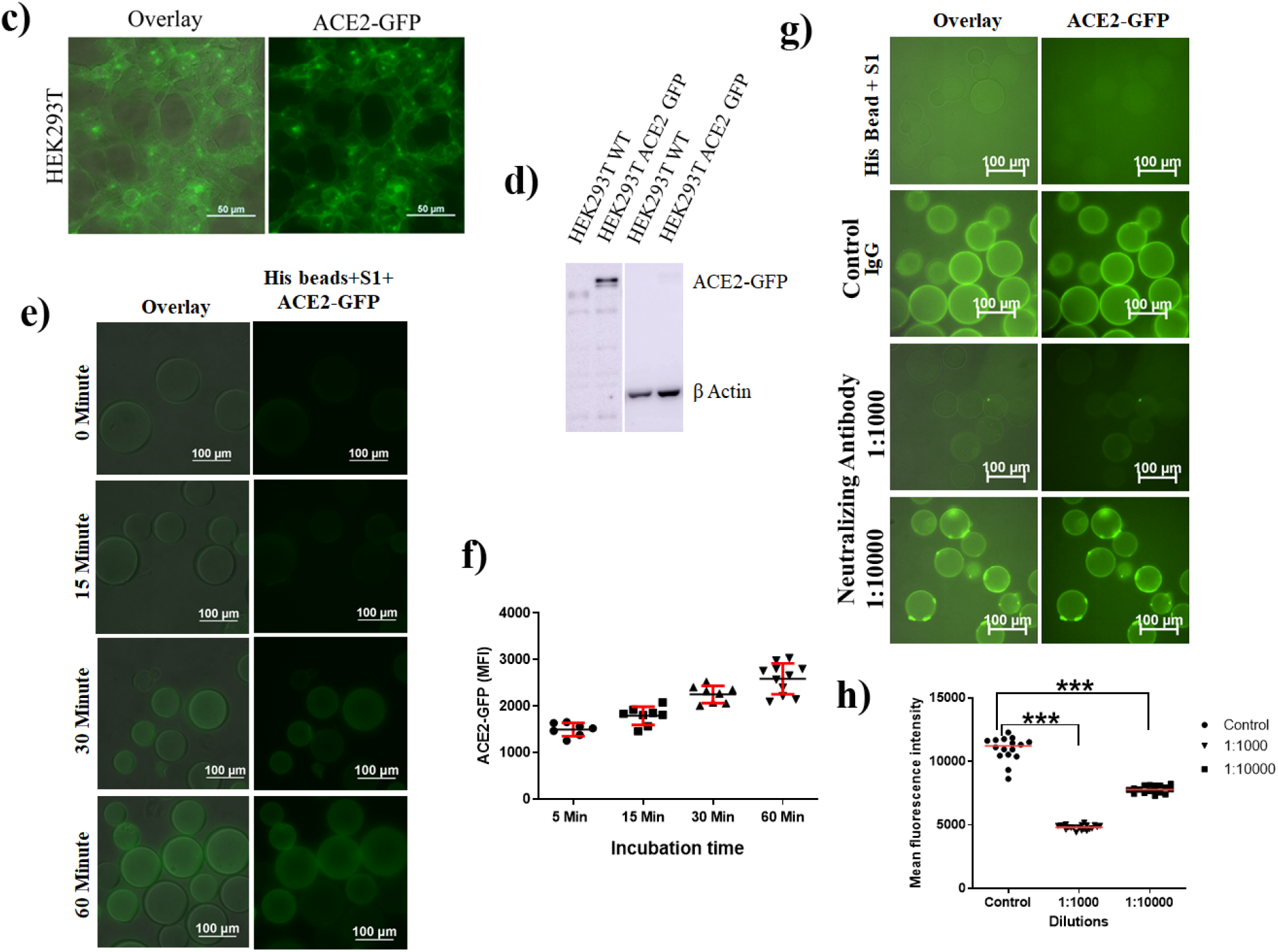
**a:** recombinant RBD domain of SARS-CoV-2 S1 protein expressed in HEK293T cells with His-tag and purified using an affinity column b: The scheme showing purified RBD-S1 protein immobilized onto agarose beads using His-tag c: HEK293T stable cells expressing ACE2-GFP, overlay, and GFP channels shown d: Immunoblotting of HEK293T cells stably expressing ACE2-GFP e: RBD-His agarose beads incubated with whole-cell extract of HEK293T-ACE2-GFP and imaged at indicated time points f: The quantification of ACE2-GFP binding on RBD-His agarose beads at indicated time points g: The RBD-His beads were incubated with the indicated dilution of neutralizing antibody for 1 hour followed by HEK293T-ACE2 GFP extract. h: The quantification of ACE2-GFP binding on RBD-His agarose beads at indicated dilution of neutralizing antibody for 1 hour. p-value calculated using student’s t-test (0.0001>***,0.001>**, 0.01>*)

Interestingly, the simple addition of the whole-cell protein extract at a concentration of 200-300 μg per ml was enough to provide an imaging-ready bead signal rendering the assay extremely easy to perform in any laboratories without protein purification. RBD beads did not bind to the purified GFP or the whole-cell extract from GFP stable cells, confirming the specificity of S1 – ACE2-GFP binding (Not shown). Initially, we employed FPLC purified RBD-His. However, it is easy to prepare RBD-bound agarose beads using commercial Ni-NTA beads directly using the whole-cell extract of HEK293T-RBD-His stable cells. Also, any simple automated microscopy configuration can be used to quantify ACE2-GFP signal on beads using 10x or 20x objectives. We have evaluated the utility of the system to test the activity of neutralizing antibodies using a commercially available neutralizing antibody from Sino Biological. The RBD beads were pre-incubated with different neutralizing antibody dilutions for 1 hour before treatment with ACE2-GFP extract. As shown in figure 1 f, compared to the isotype-specific control antibody, neutralizing antibody reduced the bead fluorescence signal of ACE2-GFP binding in a dilution-dependent manner. The mean fluorescence intensity per bead and the intensity surface plot demonstrates the bead-based system’s sensitivity and performance. Beginning from the bead preparation to assay completion, including imaging and data analysis can be completed within 3-4 hours, which is rapid compared to the gold standard assays currently used for neutralizing antibody testing. Overall, the results demonstrate the utility of a bead-based system for testing the neutralizing activity of the SARS-CoV-2 antibody using recombinant protein immobilization on solid agarose beads by fluorescence microscopy.

### GFP/OFP nanobody based immobilization approach and fluorescent bead assay

We have evaluated a different model by reversing the receptor-ligand immobilization approach. In this approach, ACE2-GFP immobilized beads were used to bind the recombinant S1-RBD in a solution that is similar to the ELISA-based surrogate neutralizing antibody testing approach commercially available consisting of non-fluorescent h ACE2 and HRP based colorimetric detection (Fenwick et al., 2021). However, we have evaluated the best-known GFP immobilization approach, GFP nanobody, the single-chain VHH antibody domain of specific binding activity against GFP and its fusion proteins (Muyldermans et al., 2009) for direct visualization. The GFP nanobody was expressed with GST-Tag in *E. coli* and immobilized on Sepharose beads as described. The simple incubation of the bacterial lysate is sufficient to create functionalized beads (Figure 2 a) and is ready for binding to the target ACE2-GFP after a simple washing procedure. Similarly, the commercially available GFP-Trap from Chromotek were alsoused. ACE2-GFP beads developed were incubated with S1-RBD-His conjugated with Cy3 as a soluble ligand. However, the signal obtained with the S1-Cy3 conjugate was not sufficient for reliable quantitation in relation to the strong GFP signal on beads (Not shown). As an easy alternate approach, recombinant fluorescent fusion of S1-OFPSpark (referred to as S1-OFP from here on) is the ligand for easy and direct binding. HEK293T cells were transfected with S1-OFP, and cells stably expressing high levels of S1-OFP were generated by FACS sorting and colony selection (Figure 2 b). As shown in figure 2 c, the whole-cell extract from S1-OFP stable cells readily binds to the ACE2-GFP beads compared to plain GFP beads. The binding intensity of S1-OFP reached a maximum in 30 to 45 minutes. Although the relative fluorescence of red over GFP was less owing to the strong GFP signal on GFP-nanobody, the neutralizing antibody pre-incubated S1-OFP showed reduced binding to the beads (data not shown), substantiating its utility to test neutralization effect.

**Figure 2.**
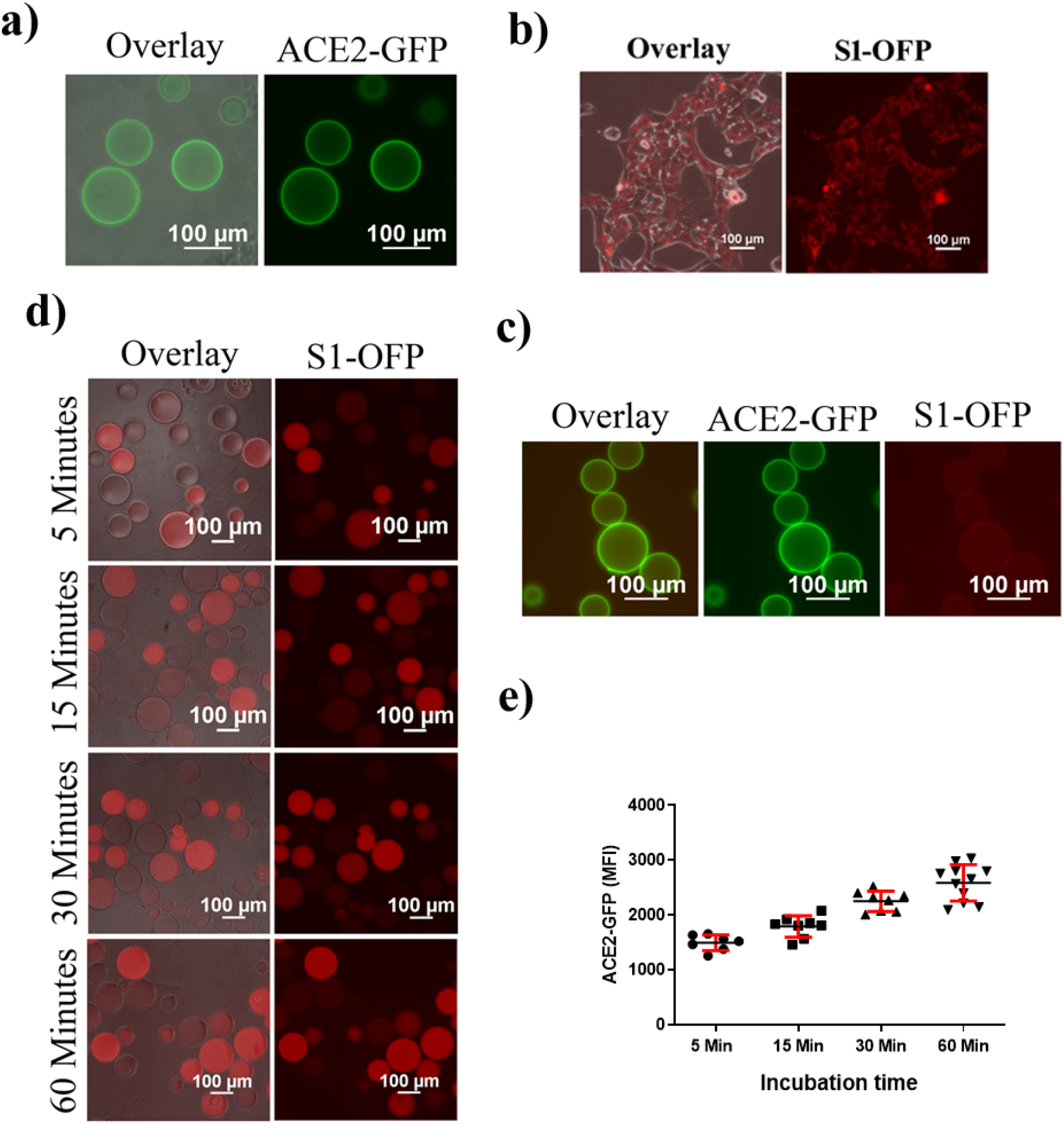
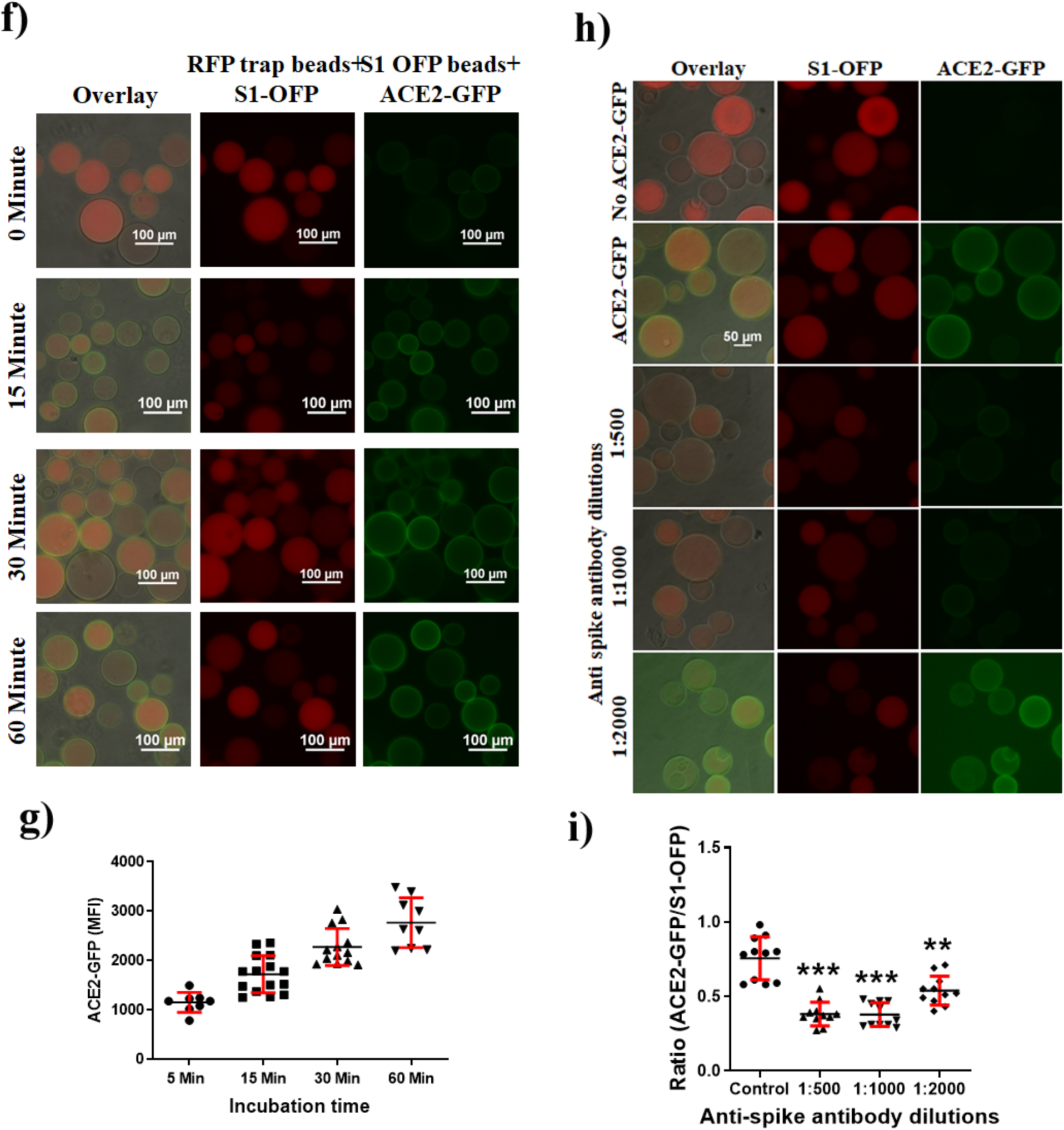
**a:** GFP nanobody with GST tag immobilized on Sepharose beads were incubated with HEK 293T ACE2-GFP lysate b: HEK 293T cells stably expressing SARS-CoV-2 S1-OFP c: whole-cell extract from HEK293T S1-OFP stable cells readily bind to the ACE2-GFP Beads d: whole-cell extract from HEK293T-S1-OFP cells at a concentration of 100 ug/ml incubated with RFP Trap beads for indicated time points, Respective overlay and RFP channels are shown e: Quantification of the S1-OFP fluorescence intensity at indicated time points are shown f: The S1-OFP beads incubated with whole cell lysate from HEK293T-ACE2-GFP cells at a concentration of 100ug/ml for indicated time points, respective overlay, GFP channel and RFP channels are shown g: Quantification of the ACE2-GFP fluorescence intensity at indicated time points are shown h: The S1-OFP beads were pre-incubated with the indicated dilution of neutralizing antibody for 1 hour followed by ACE2 GFP extract, respective overlay, GFP channel, and RFP channels are shown i: Quantification of the ACE2-GFP fluorescence intensity at indicated dilutions of neutralizing antibody are shown. p-value calculated using Two-way ANOVA with Tukey’s Post hoc hypothesis (0.0001>***,0.001>**, 0.01>*)

Considering the ease of bead pre-incubation and washing steps with the neutralizing antibody, we have immobilized the S1-OFP on beads followed by adding soluble ACE2 as the binder. Since the S1 protein is an OFP fusion, we have utilized the nanobody specific to an orange or red fluorescent protein called RFP-TRAP beads available from Chromotek for the purpose. The lysate of HEK293T cells stably expressing S1-OFP protein has been used directly for immobilizing the S1-OFP on the bead. As shown in figure 2 d, whole-cell extract from S1-OFP cells at a concentration of 100 μg/ml is robustly bound to the RFP-TRAP beads in a time-dependent manner. As shown in the figure, homogenous levels and increased orange fluorescence were noticed with RFP-tag beads within 30 minutes. The S1-OFP beads were incubated with whole cell lysate from ACE2 GFP cells at a concentration of 100 μg/ml to study the kinetics of ACE2 binding. Compared to the signal obtained from the ACE2-GFP bead, the binding of soluble ACE2-GFP on the S1-OFP bead showed a good bead signal at 60 minutes (Figure 2 e). As described, extensive washing of the beads after the binding of soluble ACE2-GFP had a negligible effect on the GFP signal suggesting direct and specific interaction of ACE2 and RBD domain of S1 protein on the beads. As shown in figure 2 e, 1:4 dilution of whole-cell extract (100mg/mL) in PBS showed a time-dependent increase in fluorescence, and by 1h, there is a significant increase in red fluorescence. Similarly, imaging after 1 hour also substantiated dilution-dependent fluorescence intensity (supplementary figure 1). To validate the utility of this system for neutralizing antibody testing, we have used the same commercial antibody. As shown in figure 2 g, the relative fluorescent intensity per bead dropped to less than 90% in neutralizing antibody pretreated bead compared to control isotype pretreated bead. The ratio of RFP/GFP served as the best readout for the determination of neutralizing activity.

### Bead assay using soluble recombinant ACE2-GFP and SARS CoV2 S1-OFP

As described above, the cells expressing the recombinant proteins were expressed stably in mammalian cells, and only high expressing clones were expanded to ensure high binding with reliable immobilization. The lysate from the stable cells can be used directly even without purification for running multiple assays with required dilutions as the beads used are functionalized in a tag-specific manner. Further, to enhance the efficiency and versatility, stable cells producing a soluble form of both ACE2 and S1 or RBD protein have been developed. The supernatant from these cells can be directly used for the easy implementation of the assay. ACE2 lacking the transmembrane domain is sufficient to render it a soluble form capable of binding to the S1 and neutralization (Verdecchia, Cavallini, Spanevello, & Angeli, 2020). HEK293T cells were stably expressed with the soluble version of ACE2 with GFP fusion. Further, the transgene’s expression in the freestyle HEK system improved soluble protein production (de Los Milagros Bassani Molinas, Beer, Hesse, Wirth, & Wagner, 2014). The image shows stable cells expressing the fusions protein and western blot confirmation of the soluble ACE2 (figure 3). The release of full-length S1-OFP from the stable cells could reflect S1 shedding or conformational change of S1 as required in host cells to assist the release of viral-RNA (Dai & Gao, 2021). Western blot revealed that the released S1-OFP reacted with the antibody against S protein, substantiating functional recombinant protein release from the stable cells developed (Figure 3 b). Subsequently, this S1-OFP immobilized on RFP-trap beads upon addition of soluble ACE2-GFP showed good binding as evident from strong bead signal (Figure 3 c). A neutralization assay performed using the commercial antibody of SARS CoV2 spike further confirmed the usefulness of the soluble recombinant proteins from the growing culture supernatant for binding, proving the approach as cost-effective and rapid one (figure 3 c).

**Figure 3.**
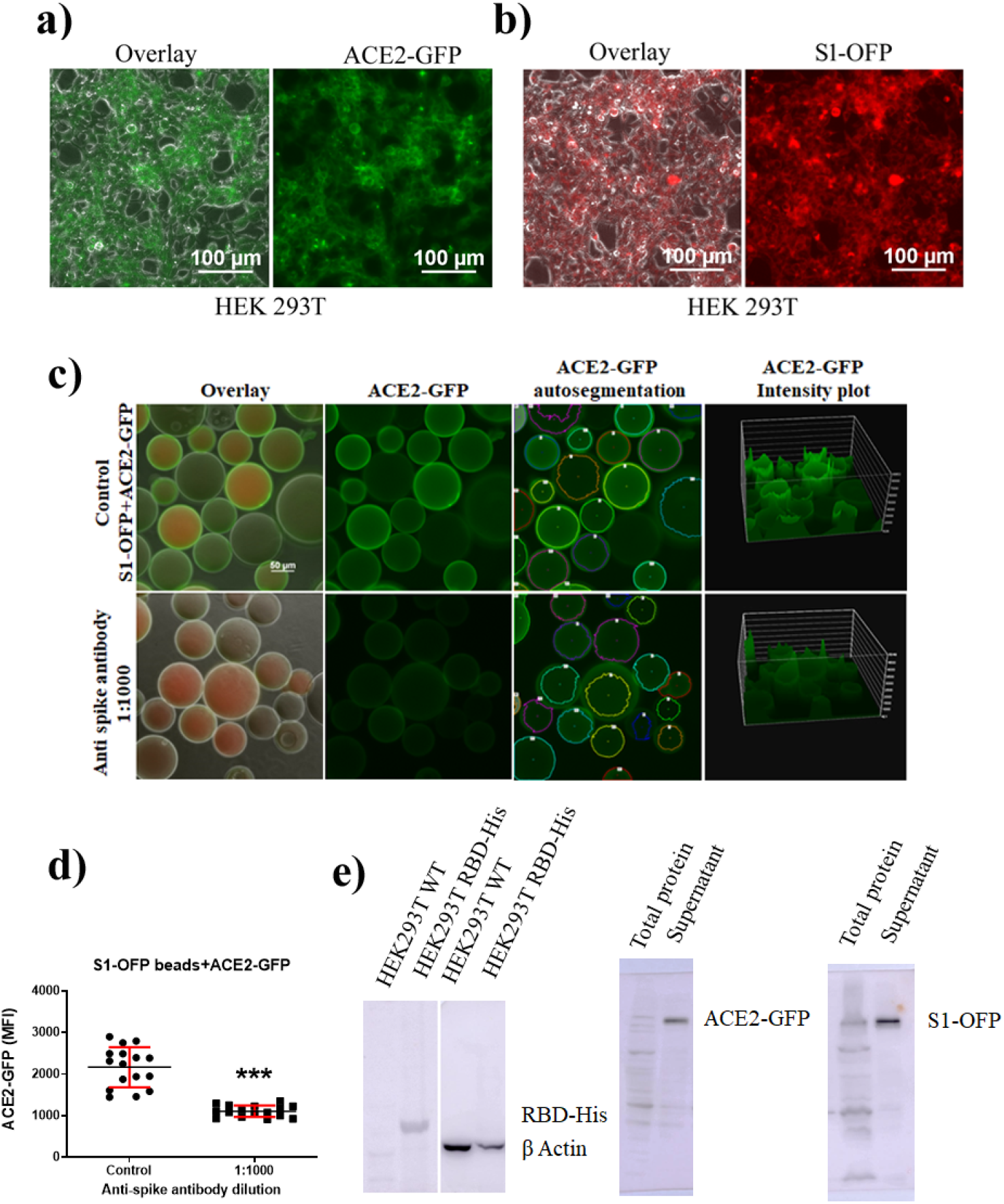
**a:** HEK293 cells stably expressing a soluble version of ACE2-GFP Fig 3 b: HEK293 cells stably expressing a soluble version of S-OFP Figure 3 c: Supernatant of soluble ACE2-GFP and S1-OFP stable cells act as ready to use reagents for HTS analysis and the quantification of the ACE2-GFP fluorescence intensity also shown Fig 3 d: Quantification of the ACE2-GFP fluorescence intensity at indicated dilution of neutralizing antibody is shown. p-value calculated using student’s t-test (0.0001>***,0.001>**, 0.01>*) Fig 3 e: Westernblotting of supernatant from HEK293T cells stably expressing soluble versions of ACE2-GFP, S1-OFP and RBD-His

### Life-time imaging confirms SARS-CoV-2-S1 – hACE2 interaction on beads

The results shown above demonstrate specific and rapid binding of soluble ACE2 on S1-OFP immobilized on beads inhibited by the neutralizing antibody. To further confirm the specificity of the S1-OFP from the stable cells and its binding to ACE2, we have used three cell lines, such as HEK293T, SiHa, and DLD1, that inherently lack hACE2 expression. HEK293T cells stably expressing HA-ACE2 were developed and used for binding studies along with vector control cells. As shown in figure 4 a, cells stably expressed with HA-ACE2 show increased specific binding to the soluble S1-OFP compared to vector control cells that failed to show any binding. Real-time confocal imaging of HEK293T-HA-ACE2 cells after exposure to culture supernatant containing S1-OFP revealed rapid binding that reached maximum intensity by 30 minutes shown in video 1. Similarly, the transient transfection of ACE2-GFP in SiHa and DLD1 further confirmed the binding specificity, as shown in supplementary figure 2.

**Figure 4.**
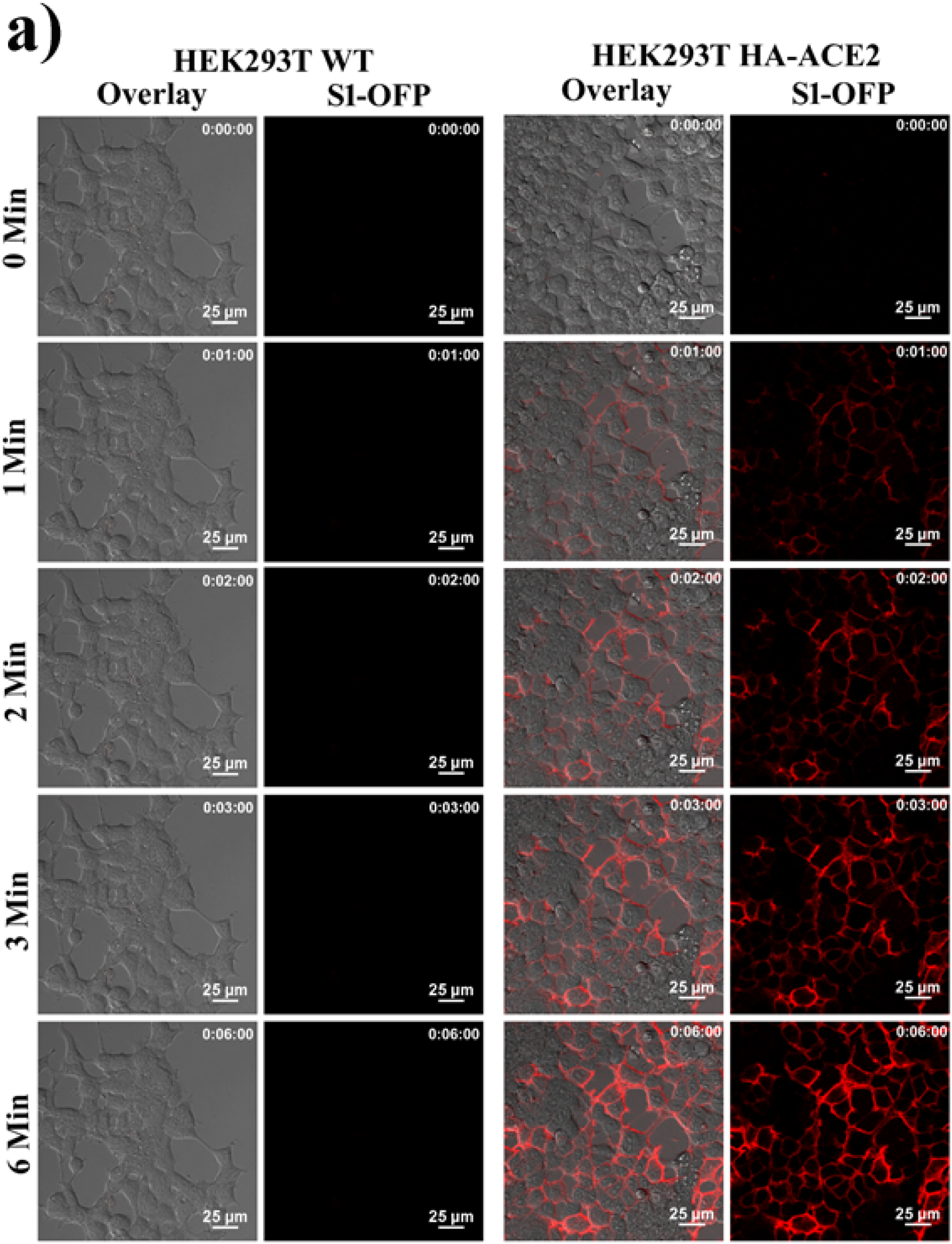
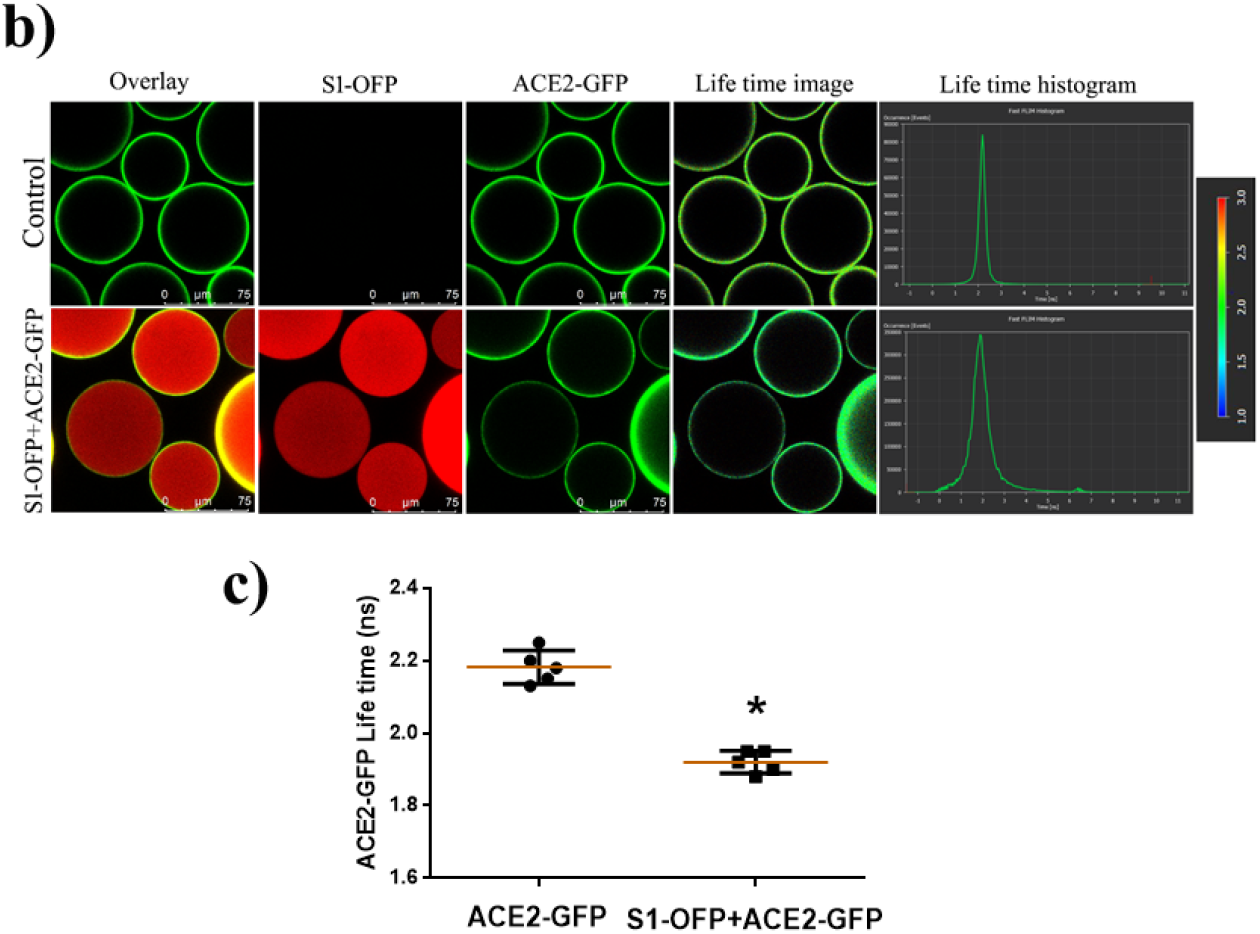
**a:** Real-time confocal imaging of HEK293T HA-ACE2 cells and HEK293T WT cells treated with S1-OFP and selected time points are shown b: Life-time imaging of ACE2-GFP bound to S1-OFP beads c: Quantification of life-time images of ACE-GFP beads and ACE2-GFP bound to S1-OFP beads. p-value calculated using student’s t-test (0.0001>***,0.001>**, 0.01>*)

The results described with beads and cells confirmed the specificity of binding of ACE2 and SARS CoV2 S1, once expressed as fluorescent protein fusion in a time and concentration-dependent manner substantiating retention of native conformation and function. GFP and orange fluorescent proteins are good FRET partners and can be used to study direct protein interactions. The most sensitive method to visualize protein-protein interaction, life-time imaging, and acceptor bleaching approach were used to confirm the interaction of ACE2 and S1-OFP on beads. As shown in figure 4 b, ACE2-GFP only beads showed a life-time of 2.2 ns that has decreased to 2.1 after binding to the S1-OFP substantiating the direct interaction of fluorescent partners on the beads. The results confirm that the bead-based assay described here using recombinant proteins retained the physiological interactions of ACE2 and S1 protein binding, the critical determinant of viral entry mechanism. The confocal acceptor bleaching further confirmed the interaction on beads. As shown in supplementary figure 3, 6-9% FRET efficiency was noticed by the acceptor bleaching method on beads.

| On-bead | In solution | On-cells<br>(HEK-293T,<br>Si-Ha,<br>DLD1) | Advantage and Disadvantages |
| --- | --- | --- | --- |
| RBD-His<br>Ni-NTA | ACE2-GFP | - | Single color read out, Good signal strength |
| ACE-2 GFP<br>(GFP-trap) | S1-RBD-Cy3 | - | Dual color readout. Low signal strength of Cy3 |
| ACE2-GFP<br>(GFP-trap) | S1-OFP | - | Dual color readout, Low signal of OFP over GFP |
| S1-OFP<br>(RFP-trap) | ACE-2GFP | - | Dual color readout, Good signal strength for both OFP and GFP |
| - | S1-OFP | HA-ACE-2 | Single color, Moderate Signal strength |

### Validation of bead assay with neutralizing antibodies and correlation with pseudovirion assay

The results shown above demonstrate the specificity of the bead-based system to report the binding of soluble ACE2-GFP to the S1-OFP bead and the nature of the interaction between the fluorescent recombinant proteins of S1 and ACE2 both on beads and cells. The bead-based system’s ability and universal application to screen diverse neutralizing antibodies was compared with the popular pseudovirion assay. To establish the correlation, Lentiviral-GFP-based pseudovirion particles pseudotyped with SARS-CoV-2-Spike protein were used. HEK293T-ACE2 stable cells were infected with the lenti-GFP/S pseudovirions, and the percentage of the infected cells counted based on the green fluorescence signal and compared with results from bead assay. We have used multiple validated commercial neutralizing antibodies, a recombinant sybody against RBD, and an in-house generated antibody in mice immunized with recombinant S1 as the antigen (Figure 5 a-d). As shown in the figures, there is a strong correlation between bead-based results and pseudovirion assay both with the mouse antibody, sybody against RBD, and three commercial antibodies tested in a concentration-dependent manner (Figure 5 e). The advantage of anti-RBD sybody is that it can be expressed in a prokaryotic system and is easy to purify the antibody using the His-affinity tag. The purification of His-tagged sybody is shown in supplementary figure 4. Overall, the results prove that the bead-based system described here is an alternate easy to perform cost-effective assay for testing the neutralizing activity of SARS-CoV-2 antibody with potential utility for the vaccine efficacy testing profiling of vaccinated population.

**Figure 5.**
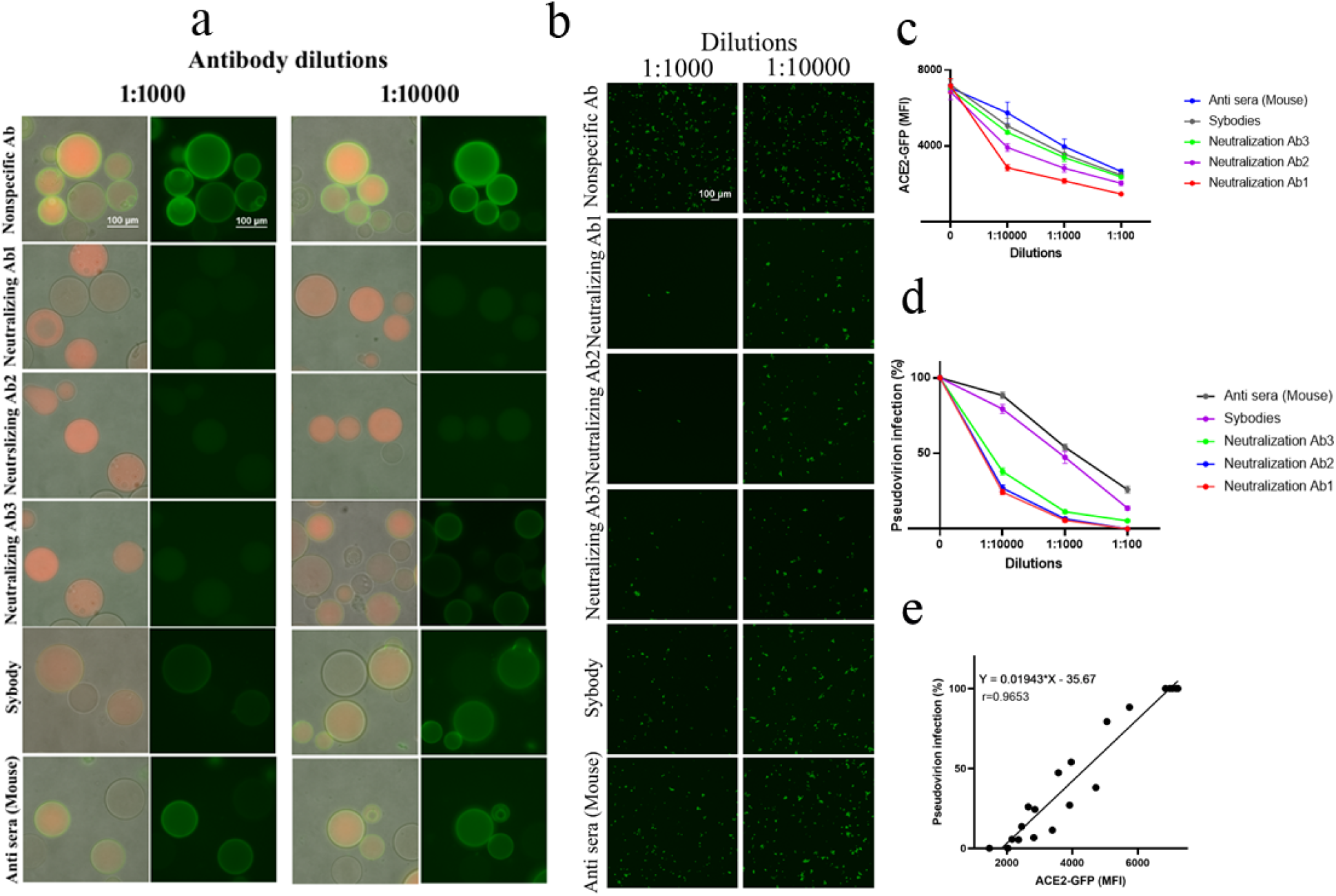
**a:** Validation of the S1-OFP-ACE2-GFP bead assay using diverse antibodies against SARS-CoV-2-S1 b: pseudovirion assay using diverse antibodies against SARS-CoV-2-S1 c: Quantification of the ACE2-GFP fluorescence intensity of S1-OFP beads incubated with indicated dilutions of antibodies d: Quantification of the pseudovirion infection (ZsGreen fluorescence intensity) incubated with indicated dilutions of antibodies e: Correlation graph showing the direct relationship between ACE2-GFP binding on S1-OFP beads with pseudovirion infection.

### Conclusion

Vaccine safety and efficacy testing are the critical determinants of the vaccine development process. Efficacy testing primarily assays for the specific neutralizing activity of antibodies generated. For SARS CoV2, neutralizing antibodies against the S1 protein or RBD domain of SARS CoV-2 is the best predictive marker of vaccine efficacy (Jiang, Hillyer, & Du, 2020). So the specific detection methods for ACE2 – S1/ RBD protein binding inhibition is of paramount importance in vaccine efficacy testing and also to understand the dynamics of protection after vaccination and infection. Here, we describe bead-based approaches to test the neutralizing antibody that specifically test the neutralizing activity of antibody to block the interaction between ACE2 – S1/RBD. The assay utilizes affinity agarose beads to bind to fluorescent proteins of SARS CoV2 S1 and human receptor h ACE2 expressed in mammalian cells. Since high expressing stable cells producing fluorescent recombinant proteins were developed, the assay protocol is rapid and can be completed within 6 hours without any laborious processing. Compared to conventional cell-based or cell-free assays, this assay is exceptionally user-friendly, economical, and scalable to multiple platforms, including automated microscopy, HTS, and flow cytometry. The assay has been used to screen diverse monoclonal neutralizing antibodies, and the results showed high concordance with the best available SARS CoV2 pseudovirion assay result. Moreover, the assay principle may be implemented for another viral-specific antibody testing, the functional utility of convalescent plasma, and comparative evaluation of diverse vaccine candidates in a rapid manner. This assay could also be used to discover viral entry blockers engaging S1-ACE2 in drug screening settings. Overall, we describe a bead-based assay platform for screening neutralizing antibodies against SARS CoV-2. The assay is expected to support and accelerate our efforts to develop newer vaccine against COVID-19 and is scalable to handle large number of samples in a rapid manner.

## Acknowledgments

This work was supported by the extramural funding from the Department of Biotechnology (BT/INF/22/SP33090/2019) and BIRAC (BT/PR40330/COT/142/16/2020).

