## Supplementary file for "A Rapid Bead-Based Assay For Screening Of SARS-CoV-2 Neutralising Antibodies"

| Company | Catalogue No | Dilutions | Purpose |
|---|---|---|---|
| Sigma | #SAB350097 8 | 1:1000 | Anti ACE2 (Western blotting) |
| Merk | #MAB8783 | 1:1000 | Anti SARS COV-2 Spike (Western blotting) |
| Sino Biological Inc | #40592-R001 | 1:100,1:1000 1:10000 | Anti SARS COV-2 Spike (Neutralizing Antibody) |
| Sino Biological Inc | #40592-MM57 | 1:100,1:1000 1:10000 | Anti SARS COV-2 Spike (Neutralizing Antibody) |
| Sino Biological Inc | #40591-MM43 | 1:100,1:1000 1:10000 | Anti SARS COV-2 Spike (Neutralizing Antibody) |

Table 1-List of antibodies and dilutions

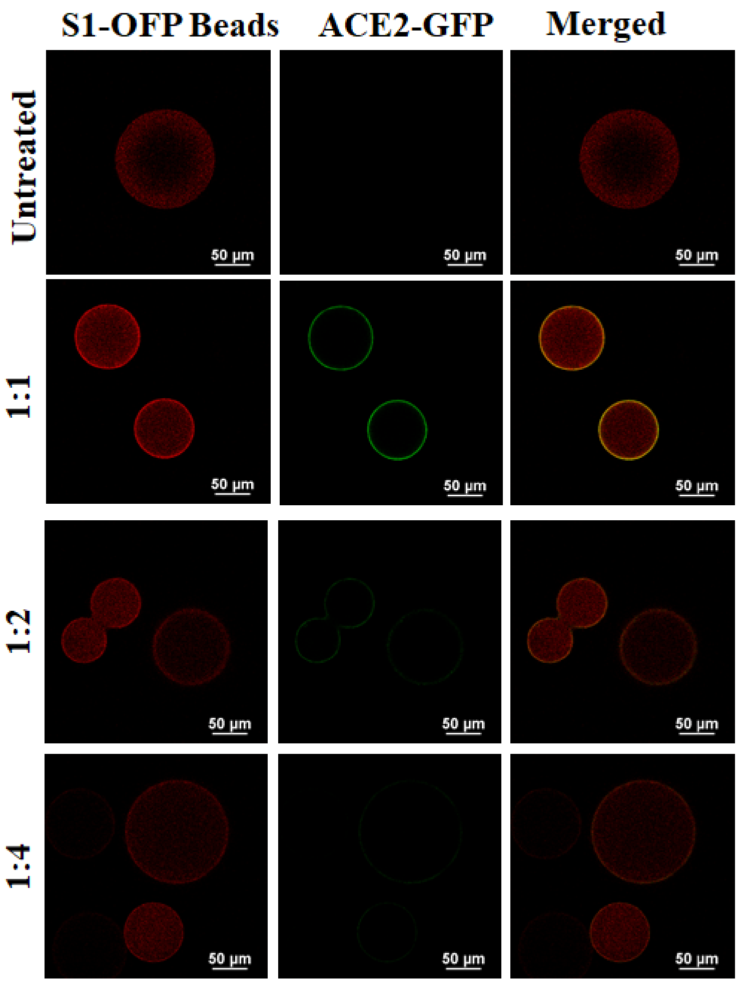


Supplementary figure 1: The S1-OFP beads were incubated with whole cell lysate from HEK293T-ACE2-GFP cells at indicated dilutions for 1hour and imaged using Confocal microscope

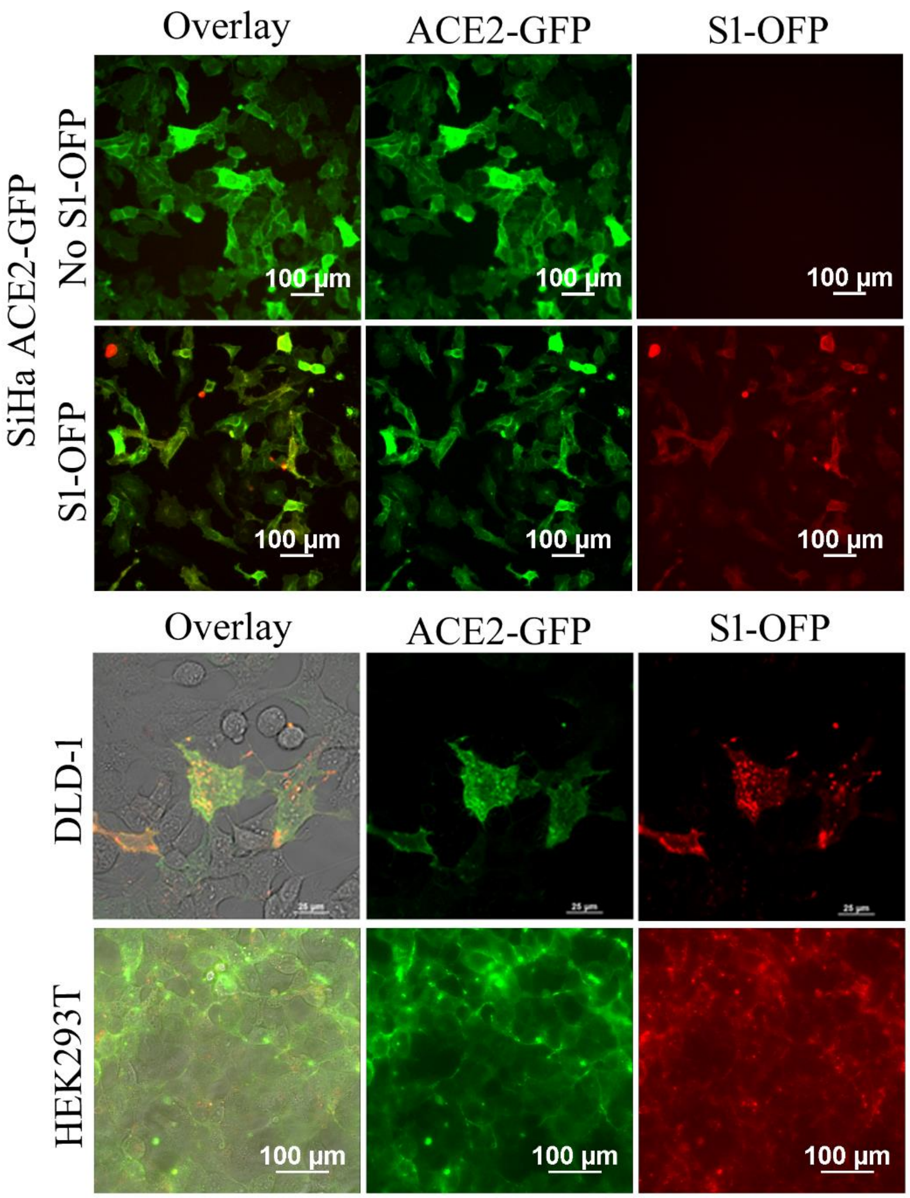


Supplementary figure 2: SiHa, DLD-1 and HEK293T ACE2-GFP stable cells incubated with S1-OFP extract for 1 hour and imaged using Widefield microscope

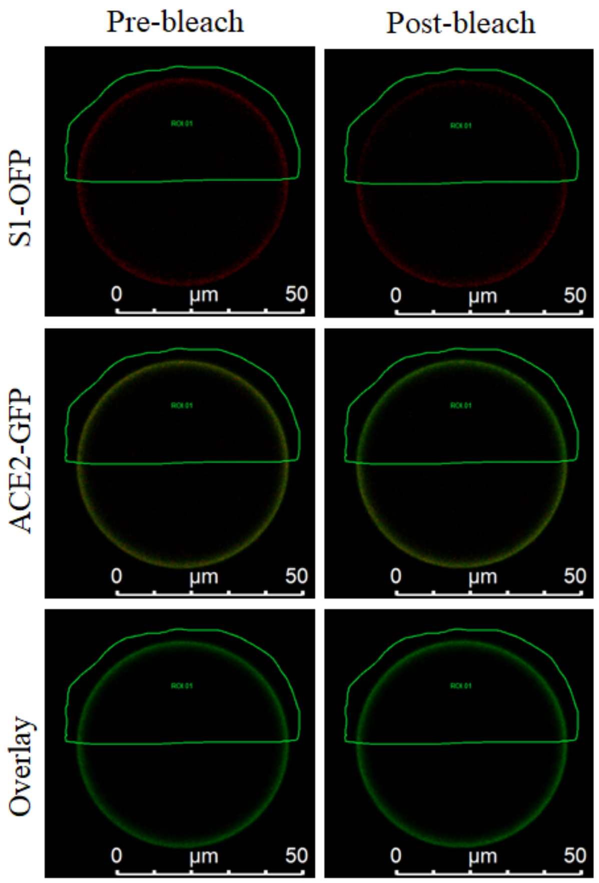


| | ROI.01 |
|---|---|
| Donor Pre | 2860.01 |
| Donor Post | 3177.65 |
| Acceptor Pre | 905.23 |
| Acceptor Post | 665.33 |
| Efficiency | 0.1 |

Supplementary figure 3: ACE2-GFP binding on S1-OFP beads confirmed using Acceptor Bleaching

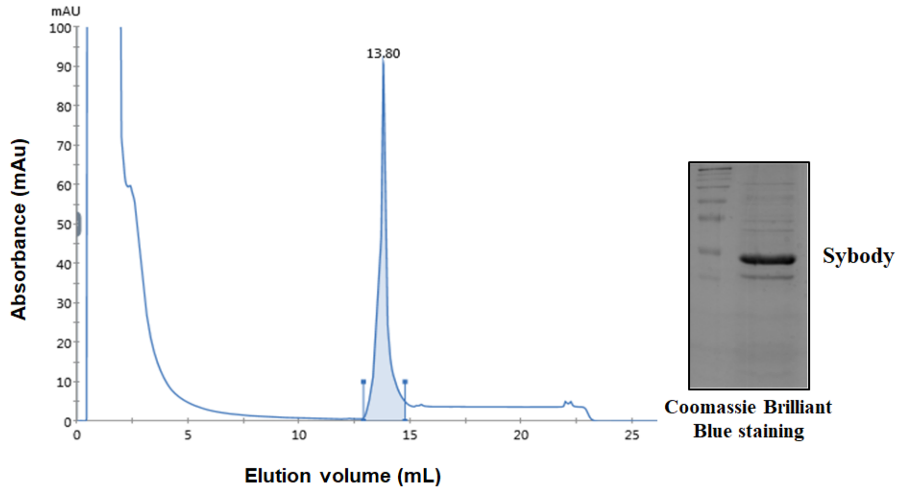


Supplementary figure 4: Sybodies Purification

Video 1: Real-time confocal imaging of HEK293T HA-ACE2 cells and HEK293T WT cells treated with S1-OFP
